# Host density dependence and environmental factors affecting laurel wilt invasion

**DOI:** 10.1101/642827

**Authors:** Robin A. Choudhury, Hong L. Er, Marc Hughes, Jason A. Smith, Grechen E. Pruett, Joshua Konkol, Randy C. Ploetz, James J. Marois, Karen A. Garrett, Ariena H.C. van Bruggen

**Author notes:** These authors contributed equally to data analysis and writing (Robin A. Choudhury, Hong L. Er). Corresponding author: R. A. Choudhury.

## Abstract

Host size, density and distribution, in addition to climate, can affect the likelihood a pathogen will invade and saturate landscapes. Laurel wilt, caused by the vector-borne forest pathogen *Raffaelea lauricola*, has devastated populations of native Lauraceae in the Southeastern US, and continues to spread. We surveyed 87 plots in six coastal islands in South Carolina, Georgia and North Florida, and one inland site (Archbold Biological Station) in South Florida for laurel wilt-affected and non-affected individual plants belonging to the genus *Persea*. The coastal island sites were surveyed once in 2008 or 2009, and the inland site was surveyed eight times from 2011 to 2013. Disease incidence per plot ranged from 0% to 96%, with mean disease incidence 45% across all sites. Disease incidence was positively correlated with trunk diameter and density of hosts with trunk diameter > 5 cm, but negatively with the degree of clustering, which was highest for small trees. A recursive partitioning model indicated that higher disease incidence was associated with moderate temperatures, wider trunk diameter, lower relative humidity, and lower wind speeds. Disease progress over time at Archbold followed a Gompertz curve, plateauing at 3% in two years. The dispersal kernel for disease incidence from a focus followed a negative exponential distribution. The number of plots with diseased trees at Archbold was similar for redbay (*P. borbonia*) and swampbay (*P. palustris*), but was lower for silkbay (*P. humilis*). Understanding how host density, size, and diversity interact with environmental effects will help guide future risk prediction efforts.

## Introduction

Invasions of pests and pathogens in naïve landscapes can rapidly and dramatically alter communities and degrade ecosystem services. Introduced pests and pathogens have devastated native forests around the world, disturbing food webs and leading to rapid environmental degradation in some cases (Akiba and Nakamura 2005; Cobb et al. 2012; Guo et al. 2019; Hughes et al. 2017; Ireland et al. 2012; Welsh et al. 2009; Wong and Daniels 2017). The density and size of hosts can significantly affect the ability of a pathogen to invade an area, as more clustered hosts can often lead to rapid transmission of disease between susceptible individuals (Anderson and May 1986; Gilbert et al. 1994; Holt et al. 2003). Environmental factors like precipitation, temperature, and relative humidity also affect the invasion of a pathogen (Anderegg et al. 2013; Shimwela et al. 2019; Welsh et al. 2009). Understanding how host and environmental factors affect pathogen invasions in forests is critical to predict where and when further spread into new regions will occur.

One of the greatest recent threats to global forest health is laurel wilt disease. Laurel wilt is caused by the ascomycete fungus *Raffaelea lauricola* T.C. Harr., Fraedrich and Aghayeva, which colonizes the vascular system of all Lauraceae hosts and can lead to rapid death in susceptible hosts (Ploetz et al. 2017). The pathogen is vectored by the redbay ambrosia beetle, *Xyleborus glabratus* Eichhoff, which carries the fungus within its mycangia (Fraedrich et al. 2008). *Xyleborus glabratus* originated in Asia (Rabaglia et al. 2006), and was first reported in the USA near the port of Savannah, Georgia, in 2002 (Fraedrich et al. 2008). The disease laurel wilt was first reported on redbay, *Persea borbonia*, in coastal forests of Georgia and South Carolina in 2003 (Hanula and Sullivan 2008). Although *R. lauricola* has been detected in *X. glabratus* in Asia (Dreaden et al. 2019; Harrington et al. 2011), laurel wilt disease has not been found in the US prior to the appearance of the *X. glabratus* (Ploetz et al. 2013). *R. lauricola* populations are quite diverse in East Asia, but the pathogen and its vector seem to be clonal in the US (Cognato et al. 2019; Dreaden et al. 2019; Hughes et al. 2017; Wuest et al. 2017). This suggests that both were introduced in a single event, possibly being carried to the port of entry in untreated packing material (Hughes et al. 2017; Ploetz et al. 2017). Since the introduction of *X. glabratus* and *R. lauricola* in the US, the disease has spread to 198 counties in nine states in the Southeastern US (USDA Forest Service 2019). Globally, laurel wilt is currently limited to the Southeastern US and parts of Asia. If this disease invades the Caribbean, or Central and South America, it would devastate both native and agricultural lauraceous hosts (Ploetz et al. 2017).

Laurel wilt affects both native and cultivated species in the Lauraceae, including redbay (*P. borbonia*), sassafras (*Sassafras albidum*), and avocado (*Persea americana*) (Ploetz et al. 2017). Redbay trees begin to show symptoms within two weeks and can die within 4-6 weeks after initial infection (Hughes et al. 2015a). Laurel wilt has decimated redbay populations and has changed the structure of plant communities throughout the Southeastern US (Goldberg and Heine 2009; Shields et al. 2011). Redbay serves as habitat and a food source for a variety of animals. It is a specific larval host for two butterfly species (Hughes et al. 2015b), and its fruits, seeds and leaves are eaten by song birds, wild turkeys, deer and black bears (Fraedrich et al. 2008). To date, an estimated 300 million redbay trees have been killed by laurel wilt (Hughes et al. 2017), roughly a third of the original native population. Other plants in the Lauraceae, such as the endangered pondberry (*Lindera melissifolia*) and gulf licaria (*Licaria trianda*), are threatened with extinction, with unknown environmental consequences (Fraedrich et al. 2011; Ploetz et al. 2017). In addition, commercial production of avocado, *P. americana* Miller, is affected in Southern Florida and threatened elsewhere (Ploetz et al. 2017; Ploetz et al. 2013).

There is limited quantitative information about the epidemiology of laurel wilt in forests, and the only previous effort to quantify the spatial expansion rate of laurel wilt in forests was completed before the disease was widely distributed (Koch and Smith 2008). Spread of laurel wilt is dependent upon redbay ambrosia beetles finding a suitable host through olfactory and visual cues (Hanula and Sullivan 2008; Mayfield and Brownie 2013). The beetles seem to prefer trees with larger stem diameters (Fraedrich et al. 2008; Mayfield and Brownie 2013). Laurel wilt host plants are interspersed with other plant species and form discrete patches in the landscape. For local disease spread, the vector must move among these patches. The densities of susceptible host plants in those patches and the distances among patches likely affect the rate of disease spread. Redbay ambrosia beetles can disperse over hundreds of meters (Hanula et al. 2016), and related scolytids are known to disperse up to tens of kilometres, depending on wind speeds and availability of suitable host materials (Barak et al. 2018; Botterweg 1982). Many environmental factors like windspeed and precipitation can favour or limit the dispersal of plant pathogens and their vectors (Narouei-Khandan et al. 2017; Shimwela et al. 2019; Shimwela et al. 2018), and prolonged environmental stress such as drought can predispose trees to bark beetle attacks and subsequent disease development (Anderegg et al. 2013; Kelsey et al. 2014; Wong and Daniels 2017).

Invasion and percolation models predict threshold host densities for increases in infection to take place (Anderson and May 1986; Holt et al. 2003). This has been observed in several different host-pathogen systems in natural plant communities (Burdon et al. 1995; Ericson et al. 1999; Jennersten et al. 1983). The relationship between infection rate and host density in natural or semi-natural plant communities can be positive (Carlsson et al. 1990; Gilbert et al. 2002; Roy 1993) or non-monotonic and unclear (Antonovics et al. 1997; Gilbert et al. 1994). The reasons for this variation can be multiple, such as the host density range, the pathogen life cycle, variation in host resistance (Garrett and Mundt 1999; Guo et al. 2019), and vector behaviour (Burdon et al. 1995; Carlsson et al. 1990; Roy 1993). Spatially explicit invasion models predict that spatial separation of hosts into patches decreases disease spread, including the spread of vector borne diseases (Caraco et al. 2001; Park et al. 2001). This decrease in spread was shown in experimental model systems (Burdon and Chilvers 1976; Neri et al. 2011), but there is little evidence for this in natural plant communities. Understanding of how laurel wilt disease spreads at a relatively small scale is still limited, and estimating a dispersal kernel may help to inform future risk models. We hypothesize that a higher density and larger size of hosts will make spread more rapid, due to an increased probability of detection by the vector, while increased clustering of host plants will decrease disease spread, due to increased difficulty for the vector to find a host across different host patches.

We studied laurel wilt dynamics in natural plant populations to determine 1) how *Persea* host attributes (including host species, diameter at breast height, host distribution and stand density) affect laurel wilt disease incidence; (2) how environmental factors like temperature, precipitation, windspeed, and relative humidity affect laurel wilt disease incidence; and (3) the temporal and spatial patterns of laurel wilt colonization in a previously unaffected area. First, laurel wilt disease distribution was analysed in six different sites across South Carolina, Georgia and Florida. Subsequently, we evaluated laurel wilt disease dynamics at another site in South-central Florida for which detailed spatio-temporal information over the course of an epidemic was available. Previous studies of laurel wilt have focused on redbay and avocado (Hanula and Sullivan 2008; Hughes et al. 2017; Ploetz et al. 2017; Shields et al. 2011). In the present work, in addition to redbay, we studied swampbay *P. palustris*, syn. *P. borbonia* var *pubescens*, and silkbay *P. humilis,* as hosts of *R. lauricola*.

## Materials and methods

### Study sites

There were seven study sites throughout the Southeastern US (Fig. 1): Edisto Island State Park and Hunting Island State Park in South Carolina; Cumberland Island National Seashore and St Catherines Island in Georgia; and Fort Clinch State Park, Fort George Island, and the Archbold Biological Station (“Archbold”) in Florida. The coastal sites are Atlantic barrier islands along the coast of the southeastern US that were created by silt and sand transported by ocean currents. The islands have sandy soils with poor water retention, high evaporation rates, and characteristic, maritime hammock vegetation comprised of live oak, *Quercus virginiana* Miller, and an understory of redbay and several other species (Goldberg and Heine 2009). Archbold is located inland on the Lake Wales Ridge, 12 km south of Lake Placid in south-central Florida. The soil is sandy with a higher organic matter content than the island soils. The vegetation is dominated by pines (*Pinus* spp.) and oaks (*Quercus* spp.), with a wide variety of understory plants, including redbay, silkbay, and swampbay. Laurel wilt had been detected at the Atlantic barrier island sites for 2-5 years prior to the survey, and was first detected approximately 0.5 km north of Archbold in early 2011.

**Fig. 1.**
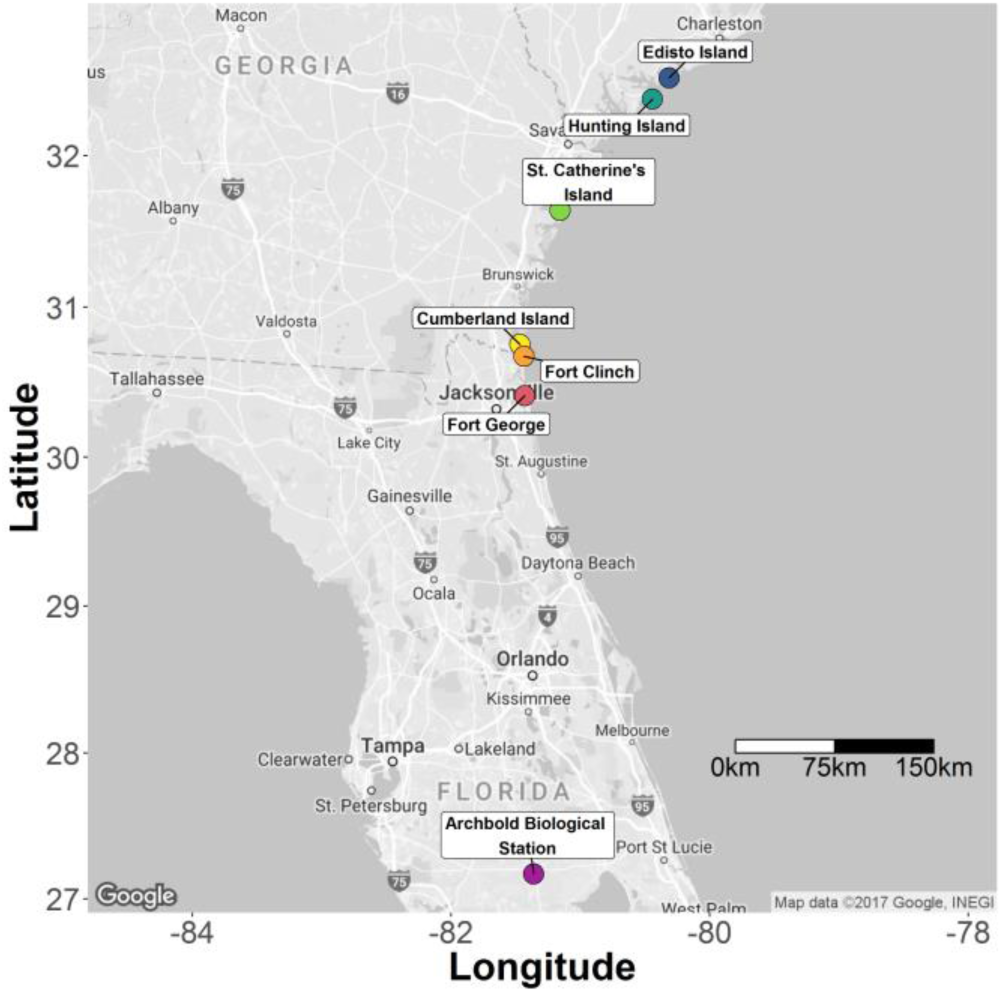
Location of study sites in South Carolina, Georgia, and Florida, in order from North to South: Edisto Beach State Park, Hunting Island State Park, St. Catherine’s Island, Cumberland Island National Seashore, Fort Clinch State Park, Fort George Island, and Archbold Biological Station.

### Survey methods

We conducted surveys in 2008 and 2009 at the six coastal sites. At least ten 32 m diameter circular plots were established at each site. Each plot had a healthy redbay tree in the center and at least one tree affected by laurel wilt. In each plot, we recorded the location and the diameter at breast height (DBH) for all symptomatic and asymptomatic redbay trees. In all cases, laurel wilt was confirmed for symptomatic host trees by the presence of discolored sapwood and subsequent isolation of *R. lauricola* (Hughes et al. 2015a). We categorized all asymptomatic trees as healthy, which likely resulted in few misclassifications due to the short latent period (approximately 4-6 weeks) of the disease. Laurel wilt incidence was calculated as the proportion of susceptible hosts affected per plot in each survey site.

We established 26 circular plots at Archbold in 2010, prior to any observations of laurel wilt disease. Each plot was 40 m in diameter and contained redbay, swampbay and/or silkbay trees for which GPS coordinates and DBH were recorded. We conducted laurel wilt surveys at intervals of one to eight months for two years (2011-2013). Laurel wilt incidence was calculated as mentioned above.

### Weather data

We collected daily weather data including precipitation, relative humidity, wind speed, and minimum and maximum temperature for 3 years prior to the survey date for all sites. In addition, we collected diurnal weather data for Archbold from an on-site weather station, accessed at www.archbold-station.org. Historical weather data for the Atlantic barrier island sites were collected from nearby airport stations (www.ncdc.noaa.gov). Weather data for Edisto Island came from Charleston Air Force Base-International (KCHS), for Hunting Island from Beaufort Marine Corps Air Station (KNBC), for Cumberland Island from McKinnon St. Simon Island Airport (KSSI), for Saint Catherines Island from Hunter Army Airfield (KSVN), for Fort George from Mayport Naval Station (KNRB), and for Fort Clinch from Jacksonville International Airport (KJAX).

### Spatial analysis

The spatial distributions of healthy and laurel wilt-affected hosts within and among plots were visualized in R version 3.4.3 (R Core Team 2013). All study sites fell in the Universal Transverse Mercator (UTM) zone 17; therefore, projected GPS coordinates in this zone were used to map the centre point of each plot. Redbay, swampbay, and silkbay host density distributions in Archbold were obtained using inverse distance weighting using the *idw* function in the *gstat* package (Pebesma and Heuvelink 2016).

### Statistical analyses

#### Clustering of host trees

The host densities per plot were visualized in GIS using R version 3.4.3 (R Core Team 2013). K-means clustering was used to assess relative clustering of trees within a plot, and silhouette indices were calculated to identify the best fitting number of clusters per plot using the *fviz_nbclust* and *kmeans* function from the *factoextra* package in the R programming environment (Kassambara and Mundt 2016). A host frequency distribution per plot was constructed for each study site. A negative binomial model was fitted to the host frequency distribution (healthy, diseased and total trees per plot) in each study site using PROC GENMOD in SAS 9.3 (SAS Institute Inc., Cary, NC). The negative binomial distribution is defined by its mean and the k parameter, 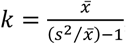, which decreases with increasing host clustering. Disease incidence was modeled as a function of the *k* parameter estimates and DBH in a generalized linear model using the *glm* function with a gaussian link function in R version 3.4.3 (R Core Team 2013).

#### Effect of host size and density on disease incidence

Mean disease incidence (the proportion of affected trees across all *Persea* spp.) for each plot was modeled as a function of mean host DBH in a generalized linear model using the *glm* function with a gaussian link function in R version 3.4.3 (R Core Team 2013). Mean disease incidence was also modeled as a function of log host density per hectare, where log transformation was selected because of a lower Aikaike Information Criterion (AIC) value than for other transformations and for the untransformed host density. The host densities for the plots were subsetted by four size categories: all trees, trees >2.5 cm DBH, trees >5 cm DBH, and trees >7.5 cm DBH. These size categories were chosen because of their relevance to forestry surveys for the region. Goodness-of-fit of the model was evaluated based on the p-value, normality of the residual distribution, and the R^2^ in regression analyses of predicted versus observed values. Pearson’s correlation coefficient and Spearman’s rank correlation coefficient were also used to estimate the strength of the relationship between the covariates.

#### Relationships between laurel wilt disease incidence and environmental and host variables

We evaluated the response of disease incidence observed in the survey plots to multiple environmental covariates. The variables used in this analysis included mean DBH, host density (the number of trees per hectare), the average minimum distance between trees inside a plot, years since first detection of disease, the mean daily high temperature for the plot (averaged over three years prior to the survey), mean daily low temperature (averaged over three years prior to the survey), mean daily high relative humidity (percentage), mean daily low relative humidity (percentage), mean daily high wind (km / h), mean daily average wind (km / h), mean daily cumulative precipitation (mm), and proportion of days with precipitation. We randomly subset 70% of our data for use as a training dataset and trained a Classification and Regression Tree (CART) model using a recursive partitioning algorithm to estimate the relative importance of the different variables with the R package *rpart* 4.1-11 (Therneau et al. 2015). To evaluate the quality of the predictions made by the CART model, we used the model to predict the disease incidence in the remaining 30% of the data, and then used a linear regression model to compare the predicted and actual values. Goodness-of-fit was evaluated as described above.

#### Epidemic development in Archbold Biological Station

The 26 plots in Archbold were surveyed every 1 to 8 months for over two years. The Gompertz model was selected to model disease incidence over time using the *gompertz* function in the R *grofit* package v. 1.1.1-1 (Kahm et al. 2010). The equation of the model was 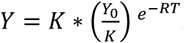, where *Y* was the observed disease incidence at time *T*. Parameters estimated were *K*, maximum disease incidence, *Y0*, the disease incidence at time zero, and *R*, the apparent infection rate. Goodness-of-fit to the model was evaluated as described above.

#### Laurel wilt dispersal gradients in Archbold Biological Station

One of the Archbold plots, coded as ABS47, had a high, early incidence of laurel wilt and was treated as the disease focus for Archbold. The negative exponential dispersal model (Campbell and Madden 1990) was used to describe the disease incidence over distance. The model was *Y* = *A * e*^−*Bx*^ (Glienke-Blanco et al. 2002), where *Y* was disease incidence at distance *x* from disease focus, *A* was disease incidence at the disease focus, and *B* was the rate of decrease in disease incidence with distance. The parameter *B* was estimated using the *fitdist* function in the R package *fitdistrplus* 1.0-9 (Delignette-Muller and Dutang 2015). Goodness-of-fit of the model was evaluated as described above.

#### Relation between laurel wilt incidence and size for the three host species in Archbold Biological Station

To test whether laurel wilt incidence in Archbold was related to host species, the difference in average DBH per plot between symptomatic and asymptomatic hosts was evaluated with Welch’s t-test separately for each host species, with corrections for uneven sampling between the different host species.

## Results

### Survey of laurel wilt-affected sites in the southeastern US

We found laurel wilt symptomatic trees at all of the surveyed sites and plots throughout our study. For the coastal sites, redbay was the only host species present, and the mean disease incidence for these sites was higher than for the inland site, ranging between 43.6% and 86.1%. The mean DBH of *Persea* trees in the coastal sites ranged from 5.5 cm to 12.4 cm, and the mean host densities of the sites ranged between 119 and 438 trees per hectare. *Persea* species are highly aggregated, based on the host densities within the surveyed plots (for an example, see Fig. S1). The weather for these sites was cooler than the inland site, and had lower variation in temperature and relative humidity. The average high wind speed tended to be higher than the inland site, although Hunting Island and Saint Catherine’s Island had lower average high wind speeds compared with Archbold. At Archbold, the hosts were identified as redbay (*P. borbonia*), swampbay (*P. palustris* or *P. borbonia* var. *pubescens*) and silkbay (*P. humilis*), with no other Lauraceae hosts present at the site. The relative frequencies of these three species differed between plots (Fig. S2). While the mean overall density of *Persea* species was 194 host individuals per hectare at Archbold, the plots ranged between 24 and 1114 host individuals per hectare. Across all of the Archbold plots, the mean DBH for redbay, swampbay and silkbay was 2.6 cm. Mean disease incidence at the end of the study across all Archbold plots was 3%. The weather at Archbold during the three years prior to the survey was more variable than the coastal sites, with greater ranges in temperature and relative humidity, while the maximum daily temperature was on average 5°C higher than that at the coastal sites.

### Effect of host size, distribution and density on disease incidence

Disease incidence was positively and strongly correlated with host DBH across all sites (Fig. 2A). Linear regression provided strong evidence that disease incidence increased with DBH (R^2^ = 0.697, p = 8.5e-24), with every centimeter increase in DBH associated with a 7% increase in disease incidence. The spatial distribution of trees with laurel wilt within individual plots was clustered (Fig. S1), and laurel wilt disease incidence increased linearly as the *k* parameter of the negative binomial distribution increased (R^2^ = 0.745, p < 2.2e-16), where increasing *k* indicates decreasing degree of clustering of laurel wilt hosts (Fig. 2B). The host DBH also increased with increasing values of the *k* parameter (R^2^=0.696, p < 2.2e-16), and thus DBH declined as clustering increased (Fig. 2C). The relationship between disease incidence and host density was not very clear when no distinction was made among host sizes (Fig. 3). When subsetting the dataset by different host sizes, disease incidence was positively correlated with host density, with Pearson’s *r* / Spearman’s (ρ) values of 0.7 / 0.56 and 0.78 / 0.65 when considering hosts larger than 5 cm and 7.5 cm, respectively. Regression of disease incidence on log-transformed density showed that density significantly affected disease incidence when considering trees larger than 7.5 cm DBH (R^2^ = 0.612, p = 2e-15), and that every log unit increase in the density of large hosts was associated with a 26.5% increase in disease incidence. Host density thresholds for disease development were not detected in this analysis.

**Fig. 2.**
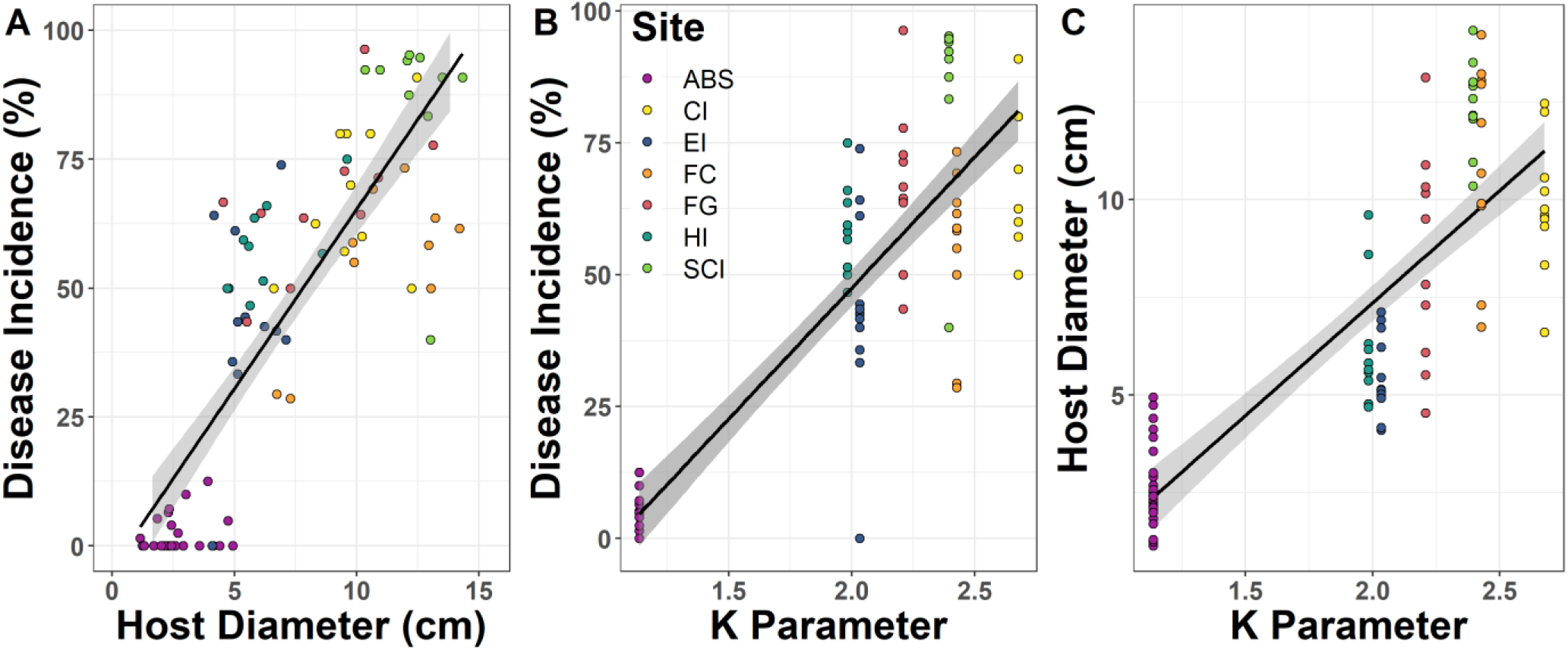
Relationships between laurel wilt disease incidence on redbay, swampbay, or silkbay and host diameter at breast height (A), between disease incidence and the *k* parameter of the negative binomial distribution (an inverse measure of clustering) (B), and between host diameter and the *k* parameter (C). Each point represents the values for a plot within a site. The smooth lines represent linear regressions, with the shaded area representing standard deviations. The regression models are: (A) y = 6.96x – 4.3 with an adjusted R^2^ value of 0.694, (B) y = 49.7x – 51.8 with an adjusted R^2^ value of 0.745, and (C) y = 5.76x – 4.2 with an adjusted R^2^ value of 0.696, respectively. The sites are Archbold Biological Station (ABS), Cumberland Island National Seashore (CI), Edisto Beach State Park (EI), Fort Clinch State Park (FC), Fort George Island (FG), Hunting Island State Park (HI), and St. Catherine’s Island (SCI).

**Fig. 3.**
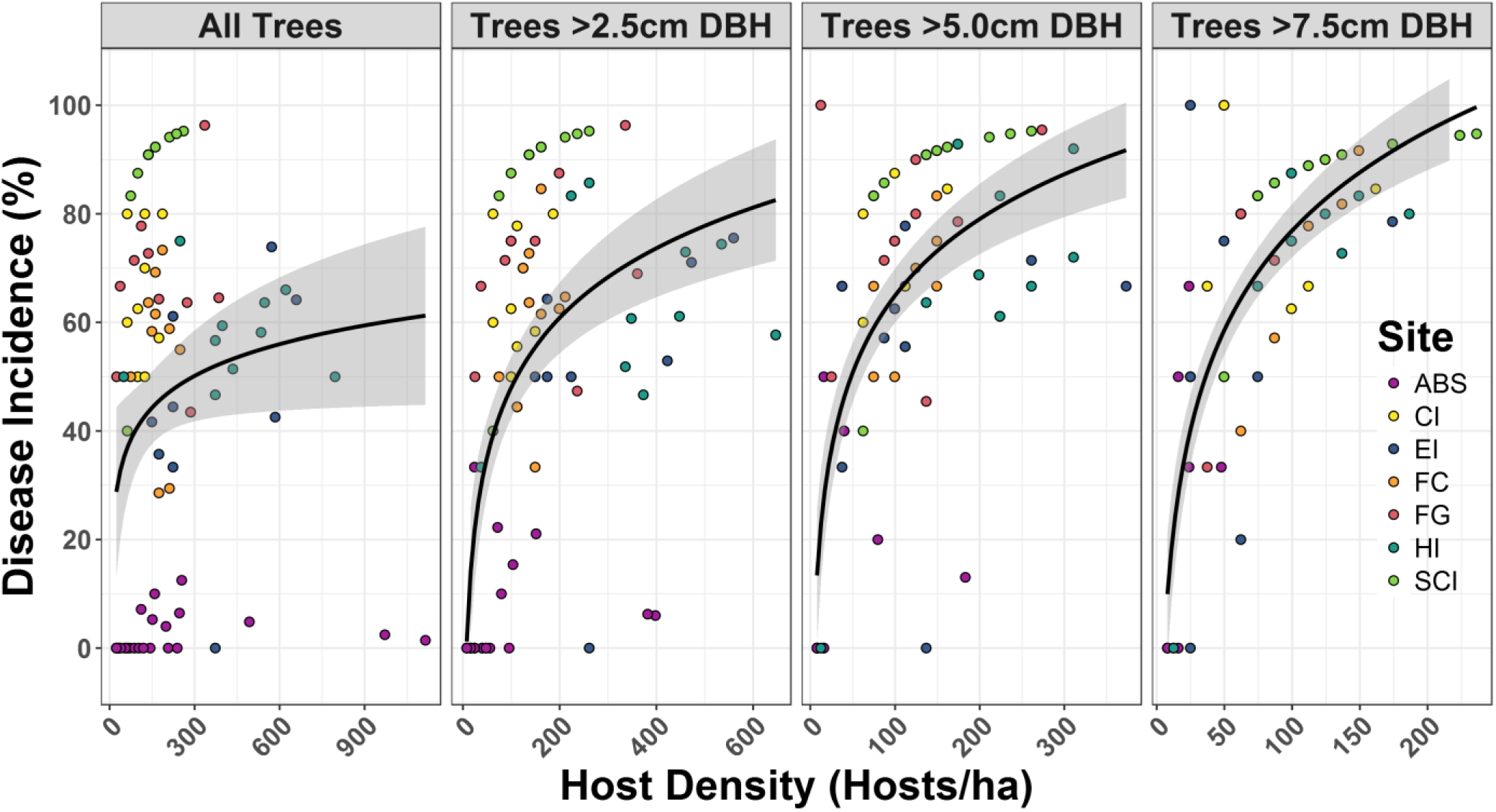
Laurel wilt disease incidence at different densities of tree hosts (redbay, swampbay and silkbay) for trees of all sizes, trees with DBH>2.5cm, trees with DBH>5.0cm and trees with DBH>7.5cm at seven sites. Smooth lines represent back-transformed data of linear regressions of disease incidence on log transformed host density per ha. The equation of regression models are: y = 8.45x + 1.98 with an adjusted R^2^ value of 0.042, y = 18.5x – 37.1 with an adjusted R^2^ value of 0.346, y = 20.4x – 28.8 with an adjusted R^2^ value of 0.482, and y = 26.5x – 44.9 with an adjusted R^2^ value of 0.606. The sites are: Archbold Biological Station (ABS), Cumberland Island National Seashore (CI), Edisto Beach State Park (EI), Fort Clinch State Park (FC), Fort George Island (FG), Hunting Island State Park (HI), and St. Catherine’s Island (SCI).

### Relationships between laurel wilt incidence and environmental and host variables

The CART model identified several environmental and host variables that were directly linked with increased levels of disease incidence when evaluated across all sites (Fig. 4A). The model suggests that higher disease incidence was linked with cooler maximum temperatures, larger host DBH, lower maximum relative humidity, and lower maximum wind speeds. Applying the CART model to the test dataset resulted in a strong correlation between predicted and actual disease incidence (Fig. 4B). The generalized linear model indicated a strong relationship between the predicted and actual disease incidence (R^2^ = 0.841, p = 1.1e-11, Fig. 4B), with a predicted slope of 0.982, suggesting a close linear correspondence between the predicted and actual disease incidence.

**Fig. 4.**
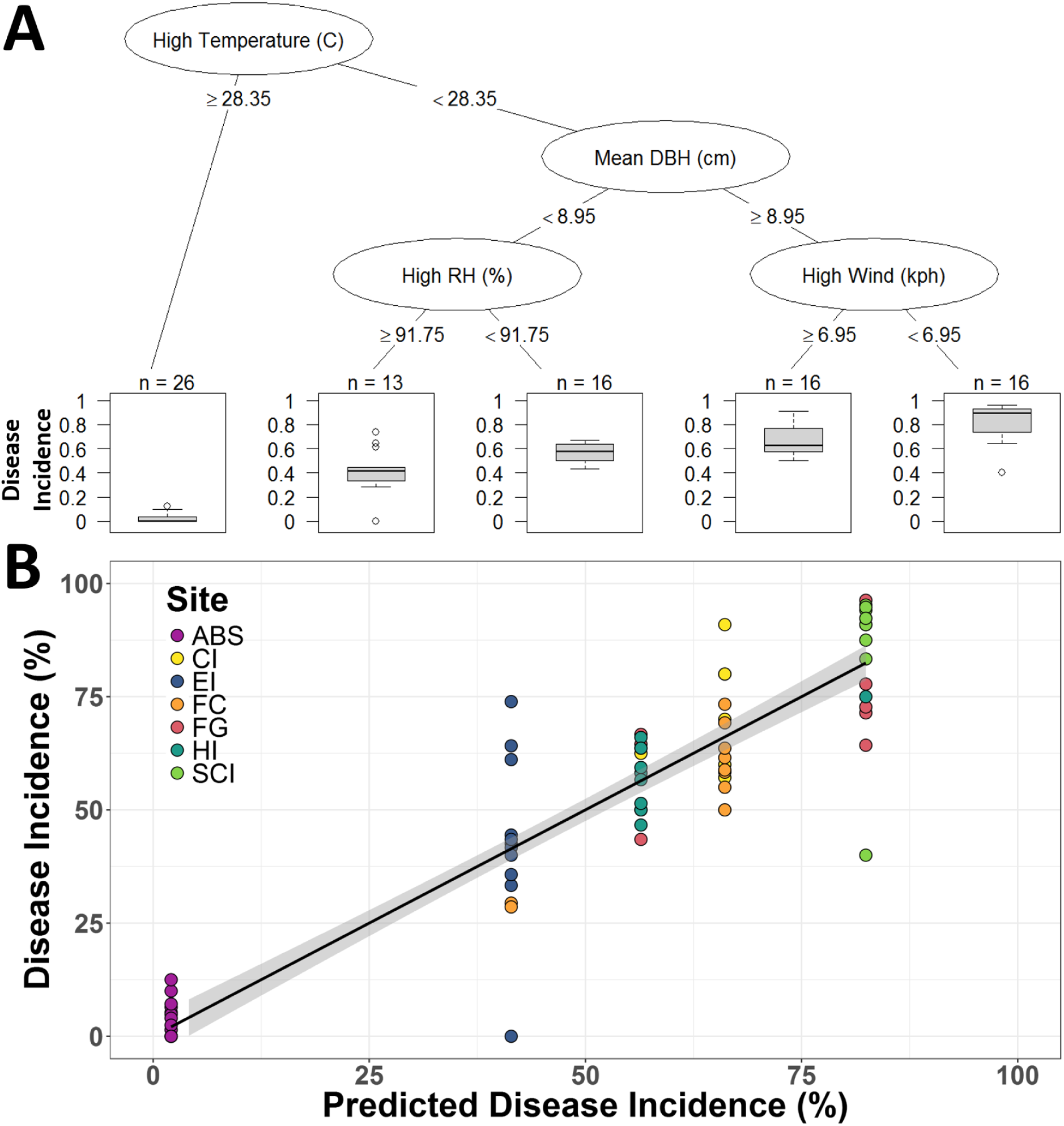
Classification and regression tree (CART) analysis of environmental and host factors related to disease incidence at different sites (A), and a regression of the predicted disease incidence (%) from each plot from the CART model and the true disease incidence values. The sites are Archbold Biological Station (ABS), Cumberland Island National Seashore (CI), Edisto Beach State Park (EI), Fort Clinch State Park (FC), Fort George Island (FG), Hunting Island State Park (HI), and St. Catherine’s Island (SCI).

### Epidemic development in Archbold Biological Station

Laurel wilt was first detected in Archbold in February, 2011, but not yet in the designated plots. This observation was taken as the zero time point for the estimation of disease progress in a Gompertz model (Fig. 5). The disease was first discovered in the designated plots in March of 2012, one year after surveys began. The model was *Disease Incidence* (%) = 3.114 *\** 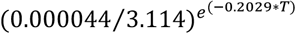. The estimated disease incidence at time zero was approximately zero and the maximum disease incidence was estimated to be 3.11%. The calculated apparent infection rate was 0.2% per month. Of the 635 trees monitored over the course of the study, only 19 (3%) developed laurel wilt symptoms by the time of the final rating in May of 2013. Ten of these 19 were redbay, six were swampbay, and three were silkbay.

**Fig. 5.**
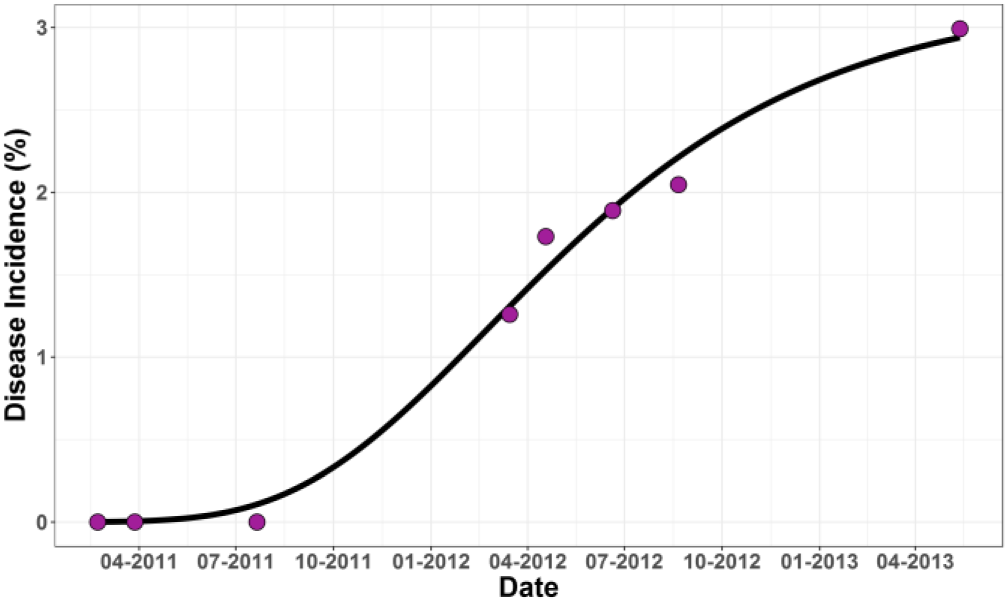
Disease progress curve of laurel wilt in Archbold Biological Station, Florida. Points represent total disease incidence across the 26 surveyed plots. The smooth line represents the Gompertz model fitted to the observed data. Equation of the Gompertz model was (DI=disease incidence, T=months): DI=3.114*(0.000044/3.114)^EXP(−0.2029*T); approximate R^2^ = 0.99

### Laurel wilt dispersal gradients in Archbold Biological Station

The initial disease outbreak in the survey area at Archbold occurred near the center of the site, in one plot coded as ABS47 (Fig. S2), with 12 plots to the north of this focus plot and 13 plots to the south. We used a negative exponential model to estimate the dispersal gradient away from this focus plot (Fig. 6). The exponential model fit the data moderately well, with R^2^ values ranging from 0.13 to 0.41. The exponential parameter had a mean value of 0.743 and a standard deviation of 0.213. The exponential parameter was slightly greater in the northern direction, suggesting that spread may have been directional over time, possibly in association with local wind patterns aiding the dispersal of the beetle vectors or due to varying temperature patterns in different parts of the Archbold Biological Station.

**Fig. 6.**
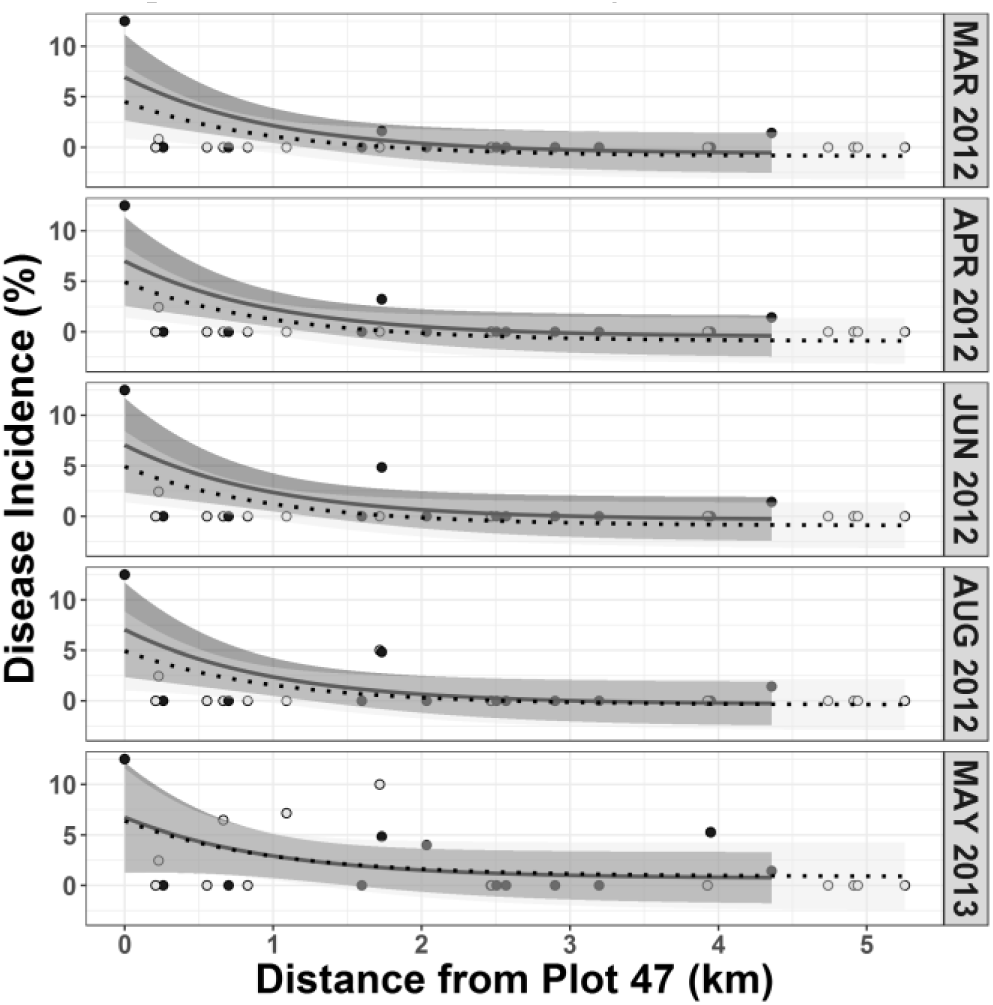
Laurel wilt dispersal gradient in the 26 surveyed plots in Archbold Biological Station North (closed points, solid lines) and South (open points, dotted lines) of disease focus ABS47 over time. Points represent observed disease incidence in each plot. Smooth lines represent exponential dispersal gradients fitted to the observed disease incidence at different distances from ABS47.

### Relation between laurel wilt and host species in Archbold Biological Station

Of the 26 plots surveyed regularly in Archbold, 10 plots only contained redbay and swampbay while 14 only contained silkbay. Two plots contained all three species. Inverse distance weighting showed that the highest densities of redbay and swampbay were in the mid-western part of Archbold, while the highest silkbay density was found in the north-eastern part of Archbold (Fig. S2). We used a Welch’s t-test to compare the DBH of laurel wilt symptomatic and asymptomatic hosts within each species. There was strong evidence for differences in the DBH of infected and uninfected plants for redbay (p < 2.2e-16, df=1251.2, t = −17.78), silkbay (p = 0.035, df = 2.04, t = - 5.11), and swampbay (p = 0.064, df=5.07, t = −2.36) (Fig. 7). The three host species were represented in a similar number of plots across the site. When considering the proportion of plots with affected individuals for each species, the ratio was similar for redbay (6 out of 12 plots) and swampbay (5 out of 11 plots), but a lower proportion of plots with silkbay (2 out of 16 plots) were affected.

**Fig. 7.**
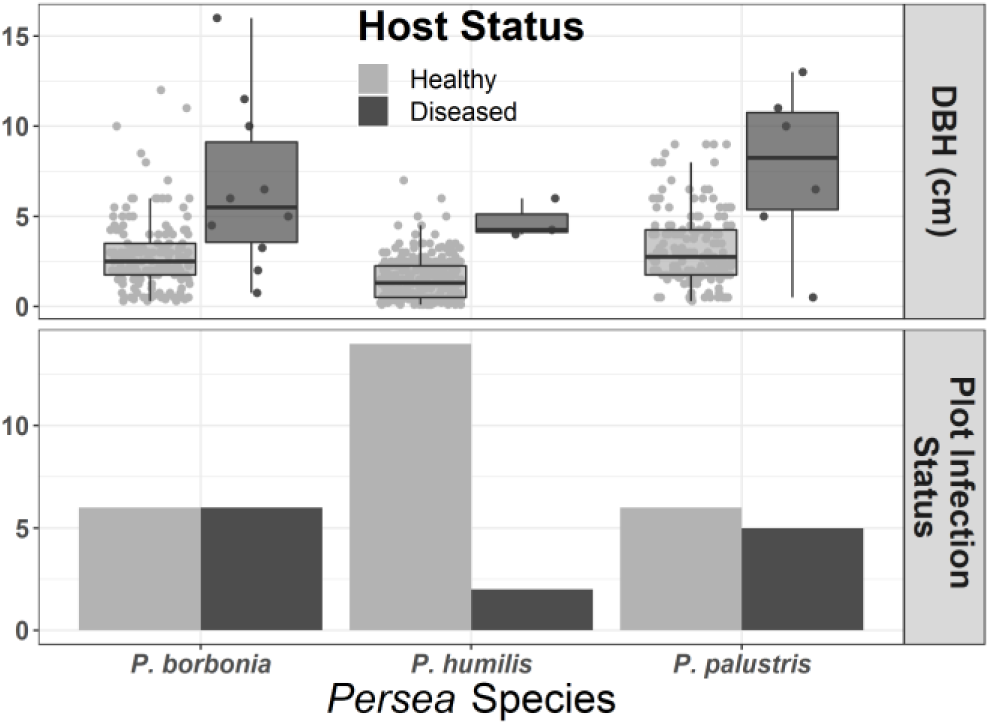
Final diameter at breast height (DBH) of healthy and laurel wilt-affected *Persea* species and the proportion of plots with affected hosts in the Archbold Biological Station site.

## Discussion

Understanding how host size, distribution and density impact the spread of invasive forest pests and pathogens is critical for predicting future outbreaks. Laurel wilt disease has devastated populations of Lauraceae in the eastern US, and the potential damage to populations in Central America, South America, and Oceania is tremendous. We found strong evidence that laurel wilt disease incidence was positively associated with increasing host size and density, but negatively with the extent of clustering. These factors may have been confounded with one another when considering landscape-level data, as observed in other host-pathogen systems in natural plant communities (Antonovics et al. 1997; Burdon et al. 1995; Carlsson et al. 1990; Ericson et al. 1999; Gilbert et al. 2002; Roy 1993). We also found that incorporating environmental variables like daily temperature, relative humidity and wind speed in a Classification and Regression Tree (CART) model improved disease incidence predictions. These results will allow us to build more accurate risk prediction models to estimate future spread of the pathogen.

Host size and age are closely linked with disease susceptibility in several plant disease pathosystems (Akiba and Nakamura 2005; Cobb et al. 2012; Gadoury et al. 2003; Reynolds and Burke 2011). Our study supports the finding that laurel wilt disease is most prevalent in trees with large stem diameter measurements (Fraedrich et al. 2008; Mayfield and Brownie 2013), and our results were consistent across multiple coastal and inland sites. While the mechanisms for age- and size-related susceptibility in other pathosystems vary from the thickness of the cambium to the emergence of susceptible plant organs, the leading hypothesis for why larger diameter trees are more frequently affected by laurel wilt is increased vector attacks. Mayfield and Brownie (2013) demonstrated that *X. glabratus* beetles home in on trees that have larger silhouette diameters. In the devastating chestnut blight pathosystem, the increased susceptibility of larger plants makes it difficult for the trees to reach sexual maturity and reproduce, limiting the species’ ability to naturally generate resistant stock through recombination (Reynolds and Burke 2011). *Persea* trees are able to reproduce to a limited extent even at small size classes, suggesting that over a long period of time some natural laurel wilt tolerance or resistance may develop within the breeding population (Hughes et al. 2015b).

Host density is frequently linked with the rate of disease increase and ultimately the total prevalence of disease in many pathosystems (Antonovics et al. 1997; Burdon et al. 1995; Carlsson et al. 1990; Gilbert et al. 2002; Roy 1993). We found that plots with higher densities of larger hosts had increased laurel wilt disease incidence, but the relationship seemed to reach a level of saturation above a certain host density. This saturation effect is similar to other pathosystems (Antonovics et al. 1997; Dobson 1990), and could be linked to various vector effects in the case of laurel wilt. High densities of large hosts occur primarily at the very beginning of the epidemic, when the numbers of ambrosia beetles are still limiting, resulting in less efficient transmission than later in the epidemic when the numbers of large trees are reduced. Alternatively, the beetle vectors may transmit the pathogen more efficiently when the host density is intermediate, but may ultimately be limited in their ability to colonize all available trees when confronted with higher densities, depending on the host distribution. Beyond natural systems, mature avocado groves represent areas of densely planted large trees, potentially attracting vectors. Maintaining avocado groves with less densely planted or younger trees may help to offset the effects of host density and size on disease incidence in agricultural settings.

Invasion models based on SIR (Susceptible - Infected - Removed) and percolation models have been used to evaluate threshold host densities at which increases in incidence take place (Anderson and May 1986; Holt et al. 2003; Ludlam et al. 2011). While infection thresholds have been demonstrated in several different pathosystems in natural plant communities (Burdon et al. 1995; Ericson et al. 1999; Jennersten et al. 1983), we did not observe clear evidence for an invasion threshold for laurel wilt in our data. This may in part be due to the growth habit of redbays, which tend to copse and form dense clusters of trees (Jones et al. 1994). It may also be due to our plot selection, where we focused on finding groups of redbays in areas already affected by laurel wilt disease. Understanding the potential role of host density in invasion thresholds will become more important as the disease begins to more heavily affect populations of Sassafras trees, which tend to grow in lower densities than coastal *Persea* species (Koch and Smith 2008). If invasion thresholds exist for this disease, the more sparse density of Sassafras may result in lower damage to Sassafras compared to redbay populations (Hughes et al. 2017). This could help limit the rate of spread of laurel wilt northwards in North America.

Understanding the role of alternative hosts is critical for predicting the severity and speed of epidemics in diverse forest settings (Cobb et al. 2012; Dobson 1990; Gilbert et al. 2002; Guo et al. 2019; Ireland et al. 2012). In some pathosystems, plant hosts can serve as super shedders, producing a prodigious amount of inoculum at the same time that they may remain asymptomatic or relatively unaffected by the disease; meanwhile, other hosts can be heavily affected by infection but produce little to no subsequent inoculum (Cobb et al. 2012). As the beetle vectors can continue to inhabit and breed in dead and dying trees, thickets of dead trees may play an important role in laurel wilt epidemics. Our study found evidence for differences in the number of plots affected by laurel wilt when invading a naïve and more diverse inland forest with higher temperatures than the more northern coastal islands surveyed. If laurel wilt disease invades naïve regions of the Caribbean and Central and South America, it will interact with previously unaffected species of Lauraceae (Ploetz et al. 2017). Some hosts (like gulf licaria, pond-spice, and pondberry) are less prevalent and only occur in small patches in the landscape (Best and Fraedrich 2018). Other hosts like the invasive camphor tree (*Cinnamomum camphora*) are relatively unaffected by the disease but can still produce inoculum (Ploetz et al. 2017). Still other hosts like avocados occur in densely planted groves surrounded by areas of native Lauraceae hosts. Understanding how laurel wilt interacts with previously unaffected species is critical to predicting how the disease will progress in new regions. Recent analyses suggest that increasing forest diversity helps to ward off pathogen and pest outbreaks (Guo et al. 2019). Many of the forests of the sub-tropics have higher diversity of Lauraceae and other tree species than those of the eastern US, which may slow down the spread of the disease.

In our study, the differences observed in laurel wilt disease incidence among redbay, swampbay, and silkbay were only partially attributable to host size, and suggest that there may be inherent differences in susceptibility of the different hosts to laurel wilt. Differences in susceptibility to laurel wilt have been shown in Lauraceae hosts under controlled and natural conditions (Fraedrich et al. 2008; Hanula and Sullivan 2008; Hughes et al. 2015a; Hughes et al. 2015b; Ploetz et al. 2017). Differences in susceptibility were quantified by differences in attraction to the redbay ambrosia beetle, population density on these hosts and/or the severity of laurel wilt symptoms. At Archbold, laurel wilt caused mortality in all three species studied (redbay, swampbay, and silkbay), and their susceptibility to laurel wilt may vary only slightly. The differences observed in the numbers of laurel wilt-affected redbays, swampbays, and silkbays may be due also to differential vector preference (Kendra et al. 2014). Understanding how different host species are susceptible or resistant to disease is important for predicting regional spread and localized extirpation of susceptible populations.

Weather and other environmental effects play a key role in predicting invasions in many pathosystems (Albrigo et al. 2005; Anderson and May 1986; Madden 1997; Shimwela et al. 2019). We used a recursive partitioning algorithm to predict host and environmental factors that were involved in laurel wilt disease incidence levels, and found that increased maximum temperatures, high relative humidities, and high wind speeds along with small stem diameters limited disease incidence within a plot. A recent paper found that found that temperatures above 26**°**C inhibited colony growth in *R. lauricola* (Zhou et al. 2018). Another recent study found that the vector *X. glabratus* is sensitive to high temperatures and increased relative humidity, with limited flight activity (Brar et al. 2012). These data suggest that laurel wilt invasion into the warmer, more humid sub-tropics may occur more slowly than previously predicted.

Laurel wilt development at Archbold was relatively slow: in slightly more than two years only 3% of the susceptible host trees had developed laurel wilt in the study plots. This is much lower than the observed incidence of laurel wilt at other sites in the present study (48-90%) and reported elsewhere over similar time frames (Fraedrich et al. 2008). The reason for the slow disease development at Archbold is unclear. Several possible explanations exist, including lower densities of single large redbay trees at Archbold compared with the coastal sites, trees that are more clustered and have smaller stem diameters, a greater diversity of vegetation types and host susceptibilities, and different weather patterns than at the coastal sites. Our study found a consistent dispersal kernel captured at a relatively large scale (∼5km). Hanula et al. (2016) monitored *X. glabratus* populations as they dispersed from a point source in a natural forest setting and found that dispersal was flat across a relatively long range (<300m). Recent work by Seo et al. (2017) suggested that the vast majority of flights completed by *X. glabratus* are short range (<10m), and only a few individuals fly over a longer range. Understanding how the vector *X. glabratus* moves across a landscape is critical for predicting how the disease will spread locally.

Predicting the spread of invasive pests and pathogens is an important step in proactive forest management. Laurel wilt has rapidly spread across the southeastern US after a single introduction event, and continues to adapt to new hosts, vectors, and environments. The disease has already wreaked havoc on native populations of redbay, threatens avocado production in Florida, and continues to impact new hosts as it spreads west and north. The impact of the disease on the Caribbean and Central and South America could potentially be devastating, disrupting diverse Lauraceae forests, although it is still unclear if the high temperatures would temper epidemic development. A better understanding of the host and environmental factors that contribute to the rapid spread of the disease is critical for understanding this ecological threat.

## Acknowledgements

This work was funded partially by USDA grants 2008-31135-19505, 2009-51181-05915, and 2015-51181-24257. We thank Drs. Gurpreet Brar, Christopher Buddenhagen, Andy Garcia, Jim Hanula, Jiri Hulcr, Russ Mizell, Roberta Pickert, Youliang Qiu, and Yanru Xing for helpful discussions and input.

---

**Fig. S1.**
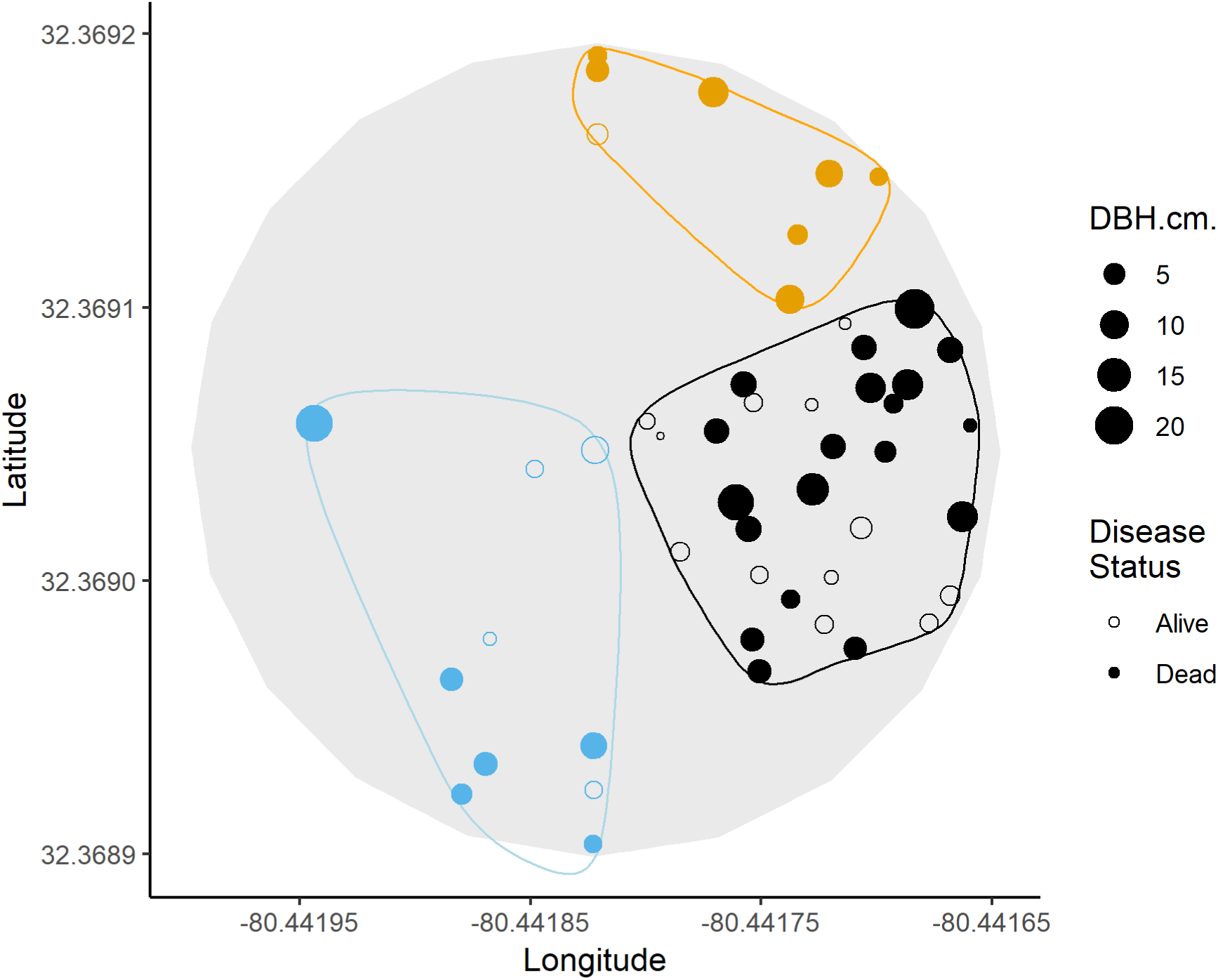
Spatial distribution of redbay (*P. borbonia*) trees in one of the plots on Hunting Island, South Carolina. The visual representation was obtained by k-means clustering in R using the factoextra package. Dead and alive trees are indicated by closed and open circles. Stem diameters at breast height are indicated by the size of the circles.

**Fig. S2.**
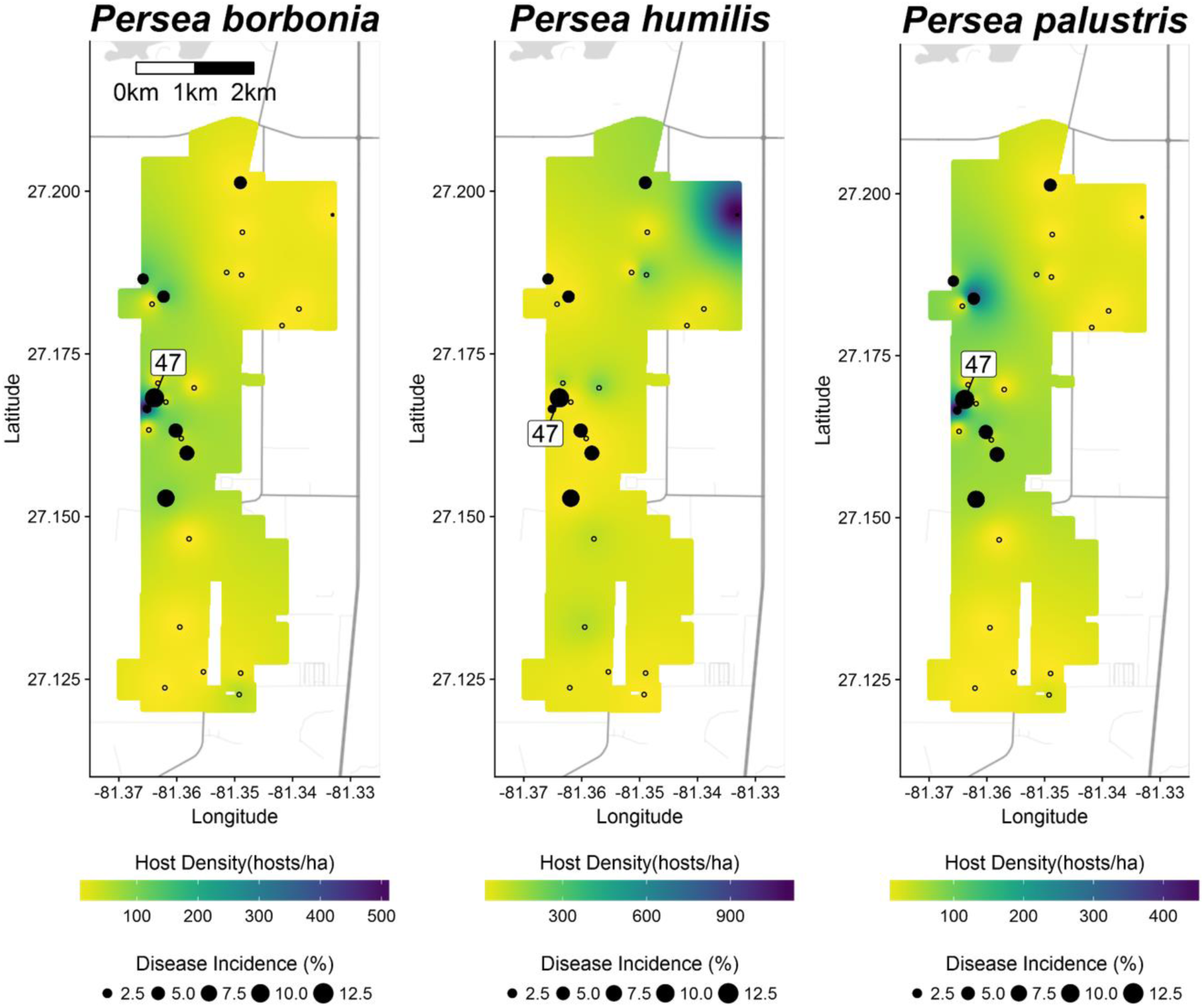
Distribution of redbay (*Persea borbonia*), silkbay (*P. humilis*), and swampbay (*P. palustris*) densities based on inverse distance weighting and total laurel wilt disease incidence for hosts (points) at Archbold Biological Station in May 2013. Empty points are plots that had no detectible infected plants.

## References

Akiba, M., and Nakamura, K. 2005. Susceptibility of adult trees of the endangered species *Pinus armandii* var. amamiana to pine wilt disease in the field. Journal of Forest Research 10:3–7.

Albrigo, L. G., Buker, R. S., Burns, J. K., Castle, W. S., Futch, S., McCoy, C. W., Muraro, R. P., Rogers, M. E., Syvertsen, J. P., and Timmer, L. 2005. The impact of four hurricanes in 2004 on the Florida citrus industry: Experiences and lessons learned. Pages 66–74 in: Proc. Fla. State Hort. Soc.

Anderegg, W. R., Kane, J. M., and Anderegg, L. D. 2013. Consequences of widespread tree mortality triggered by drought and temperature stress. Nature Climate Change 3:30.

Anderson, R. M., and May, R. M. 1986. The invasion, persistence and spread of infectious diseases within animal and plant communities. Philosophical Transactions of the Royal Society of London. B, Biological Sciences 314:533–570.

Antonovics, J., Thrall, P. H., and Jarosz, A. M. 1997. Genetics and the spatial ecology of species interactions: the Silene-Ustilago system. Princeton University Press, Princeton, NJ.

Barak, A., McGrevy, D., and Tokaya, G. 2018. Dispersal and re-capture of marked, overwintering *Tomicus piniperda* (Coleoptera: Scolytidae) from Scotch pine bolts. The Great Lakes Entomologist 33:1.

Best, G. S., and Fraedrich, S. W. 2018. An assessment of the potential impact of laurel wilt on clonal populations of *Lindera melissifolia* (Pondberry). Southeastern Naturalist 17:616–629.

Botterweg, P. 1982. Dispersal and flight behaviour of the spruce bark beetle *Ips typographus* in relation to sex, size and fat content. Zeitschrift für angewandte Entomologie 94:466–489.

Brar, G. S., Capinera, J. L., McLean, S., Kendra, P. E., Ploetz, R. C., and Peña, J. E. 2012. Effect of trap size, trap height and age of lure on sampling *Xyleborus glabratus* (Coleoptera: Curculionidae: Scolytinae), and its flight periodicity and seasonality. Florida Entomologist:1003-1011.

Burdon, J., and Chilvers, G. 1976. The effect of clumped planting patterns on epidemics of damping-off disease in cress seedlings. Oecologia 23:17–29.

Burdon, J., Ericson, L., and Muller, W. 1995. Temporal and spatial changes in a metapopulation of the rust pathogen *Triphragmium ulmariae* and its host, *Filipendula ulmaria*. Journal of Ecology:979–989.

Campbell, C. L., and Madden, L. V. 1990. Introduction to plant disease epidemiology. John Wiley & Sons.

Caraco, T., Duryea, M. C., Glavanakov, S., Maniatty, W., and Szymanski, B. K. 2001. Host spatial heterogeneity and the spread of vector-borne infection. Theoretical Population Biology 59:185–206.

Carlsson, U., Elmqvist, T., Wennstrom, A., and Ericson, L. 1990. Infection by pathogens and population age of host plants. Journal of Ecology:1094–1105.

Cobb, R. C., Filipe, J. A., Meentemeyer, R. K., Gilligan, C. A., and Rizzo, D. M. 2012. Ecosystem transformation by emerging infectious disease: loss of large tanoak from California forests. Journal of Ecology 100:712–722.

Cognato, A. I., Smith, S. M., Li, Y., Pham, T. H., and Hulcr, J. 2019. Genetic variability among *Xyleborus glabratus* populations native to southeast Asia (Coleoptera: Curculionidae: Scolytinae: Xyleborini) and the description of two related species. Journal of Economic Entomology.

Delignette-Muller, M. L., and Dutang, C. 2015. fitdistrplus: An R package for fitting distributions. Journal of Statistical Software 64:1–34.

Dobson, A. 1990. Models for multi-species parasite-host communities. Pages 261–288 in: Parasite Communities: Patterns and Processes. Springer.

Dreaden, T., Hughes, M., Ploetz, R., Black, A., and Smith, J. 2019. Genetic analyses of the laurel wilt pathogen, *Raffaelea lauricola*, in Asia provide clues on the source of the clone that is responsible for the current USA epidemic. Forests 10:37.

Ericson, L., Burdon, J., and Müller, W. 1999. Spatial and temporal dynamics of epidemics of the rust fungus *Uromyces valerianae* on populations of its host *Valeriana salina*. Journal of Ecology 87:649–658.

Fraedrich, S. W., Harrington, T. C., Rabaglia, R. J., Ulyshen, M. D., Mayfield, A. E., Hanula, J. L., Eickwort, J. M., and Miller, D. R. 2008. A fungal symbiont of the redbay ambrosia beetle causes a lethal wilt in redbay and other lauraceae in the southeastern United States. Plant Disease 92:215–224.

Fraedrich, S. W., Harrington, T. C., Bates, C. A., Johnson, J., Reid, L. S., Best, G. S., Leininger, T. D., and Hawkins, T. S. 2011. Susceptibility to laurel wilt and disease incidence in two rare plant species, pondberry and pondspice. Plant Disease 95:1056–1062.

Gadoury, D. M., Seem, R. C., Ficke, A., and Wilcox, W. F. 2003. Ontogenic resistance to powdery mildew in grape berries. Phytopathology 93:547–555.

Garrett, K., and Mundt, C. 1999. Epidemiology in mixed host populations. Phytopathology 89:984–990.

Gilbert, G., Foster, R., and Hubbell, S. 1994. Density and distance-to-adult effects of a canker disease of trees in a moist tropical forest. Oecologia 98:100–108.

Gilbert, G. S., Ferrer, A., and Carranza, J. 2002. Polypore fungal diversity and host density in a moist tropical forest. Biodiversity & Conservation 11:947–957.

Glienke-Blanco, C., Aguilar-Vildoso, C. I., Vieira, M. L. C., Barroso, P. A. V., and Azevedo, J. L. 2002. Genetic variability in the endophytic fungus *Guignardia citricarpa* isolated from citrus plants. Genetics and Molecular Biology 25:251–255.

Goldberg, N., and Heine, J. 2009. A comparison of arborescent vegetation pre-(1983) and post-(2008) outbreak of the invasive species the Asian ambrosia beetle Xyleborus glabratus in a Florida maritime hammock. Plant Ecology & Diversity 2:77–83.

Guo, Q., Fei, S., Potter, K. M., Liebhold, A. M., and Wen, J. 2019. Tree diversity regulates forest pest invasion. Proceedings of the National Academy of Sciences:201821039.

Hanula, J. L., and Sullivan, B. 2008. Manuka oil and phoebe oil are attractive baits for *Xyleborus glabratus* (Coleoptera Scolytinae), the vector of laurel wilt. Environmental Entomology 37:1403–1409.

Hanula, J. L., Mayfield, A. E., Reid, L. S., and Horn, S. 2016. Influence of trap distance from a source population and multiple traps on captures and attack densities of the redbay ambrosia beetle (Coleoptera: Curculionidae: Scolytinae). Journal of Economic Entomology 109:1196–1204.

Harrington, T. C., Yun, H. Y., Lu, S.-S., Goto, H., Aghayeva, D. N., and Fraedrich, S. W. 2011. Isolations from the redbay ambrosia beetle, *Xyleborus glabratus*, confirm that the laurel wilt pathogen, *Raffaelea lauricola*, originated in Asia. Mycologia 103:1028–1036.

Holt, R. D., Dobson, A. P., Begon, M., Bowers, R. G., and Schauber, E. M. 2003. Parasite establishment in host communities. Ecology Letters 6:837–842.

Hughes, M., Inch, S., Ploetz, R., Er, H., van Bruggen, A., and Smith, J. 2015a. Responses of swamp bay, *Persea palustris*, and avocado, *Persea americana*, to various concentrations of the laurel wilt pathogen, *Raffaelea lauricola*. Forest Pathology 45:111–119.

Hughes, M., Riggins, J., Koch, F., Cognato, A., Anderson, C., Formby, J., Dreaden, T., Ploetz, R., and Smith, J. 2017. No rest for the laurels: symbiotic invaders cause unprecedented damage to southern USA forests. Biological Invasions 19:2143–2157.

Hughes, M. A., Smith, J., Ploetz, R., Kendra, P., Mayfield, A., Hanula, J., Hulcr, J., Stelinski, L., Cameron, S., and Riggins, J. 2015b. Recovery plan for laurel wilt on redbay and other forest species caused by *Raffaelea lauricola* and disseminated by *Xyleborus glabratus*. Plant Health Progress 16:173–210.

Ireland, K., Hüberli, D., Dell, B., Smith, I., Rizzo, D., and Hardy, G. S. J. 2012. Potential susceptibility of Australian native plant species to branch dieback and bole canker diseases caused by *Phytophthora ramorum*. Plant Pathology 61:234–246.

Jennersten, O., Nilsson, S. G., and Wästljung, U. 1983. Local plant populations as ecological islands: the infection of *Viscaria vulgaris* by the fungus *Ustilago violacea*. Oikos:391–395.

Jones, R. H., Sharitz, R. R., James, S. M., and Dixon, P. M. 1994. Tree population dynamics in seven South Carolina mixed-species forests. Bulletin of the Torrey Botanical Club:360–368.

Kahm, M., Hasenbrink, G., Lichtenberg-Fraté, H., Ludwig, J., and Kschischo, M. 2010. grofit: fitting biological growth curves with R. Journal of Statistical Software 33:1–21.

Kassambara, A., and Mundt, F. 2016. Factoextra: extract and visualize the results of multivariate data analyses. R package version 1:2016.

Kelsey, R. G., Gallego, D., Sánchez-García, F., and Pajares, J. 2014. Ethanol accumulation during severe drought may signal tree vulnerability to detection and attack by bark beetles. Canadian Journal of Forest Research 44:554–561.

Kendra, P. E., Montgomery, W. S., Niogret, J., Pruett, G. E., Mayfield, A. E., III, MacKenzie, M., Deyrup, M. A., Bauchan, G. R., Ploetz, R. C., and Epsky, N. D. 2014. North American Lauraceae: terpenoid emissions, relative attraction and boring preferences of redbay ambrosia beetle, *Xyleborus glabratus* (Coleoptera: Curculionidae: Scolytinae). PLOS ONE 9:e102086.

Koch, F. H., and Smith, W. D. 2008. Spatio-temporal analysis of *Xyleborus glabratus* (Coleoptera: Circulionidae: Scolytinae) invasion in Eastern U.S. forests. Environmental Entomology 37:442–452.

Ludlam, J. J., Gibson, G. J., Otten, W., and Gilligan, C. A. 2011. Applications of percolation theory to fungal spread with synergy. Journal of The Royal Society Interface 9:949–956.

Madden, L. 1997. Effects of rain on splash dispersal of fungal pathogens. Canadian Journal of Plant Pathology 19:225–230.

Mayfield, A. E., and Brownie, C. 2013. The redbay ambrosia beetle (Coleoptera: Curculionidae: Scolytinae) uses stem silhouette diameter as a visual host-finding cue. Environmental Entomology 42:743–750.

Narouei-Khandan, H., Harmon, C., Harmon, P., Olmstead, J., Zelenev, V., van der Werf, W., Worner, S. P., Senay, S., and van Bruggen, A. 2017. Potential global and regional geographic distribution of Phomopsis vaccinii on Vaccinium species projected by two species distribution models. European Journal of Plant Pathology 148:919–930.

Neri, F. M., Bates, A., Füchtbauer, W. S., Pérez-Reche, F. J., Taraskin, S. N., Otten, W., Bailey, D. J., and Gilligan, C. A. 2011. The effect of heterogeneity on invasion in spatial epidemics: from theory to experimental evidence in a model system. PLoS computational biology 7:e1002174.

Park, A. W., Gubbins, S., and Gilligan, C. A. 2001. Invasion and persistence of plant parasites in a spatially structured host population. Oikos 94:162–174.

Pebesma, E., and Heuvelink, G. 2016. Spatio-temporal interpolation using gstat. RFID Journal 8:204–218.

Ploetz, R., Kendra, P., Choudhury, R., Rollins, J., Campbell, A., Garrett, K., Hughes, M., and Dreaden, T. 2017. Laurel wilt in natural and agricultural ecosystems: understanding the drivers and scales of complex pathosystems. Forests 8:48.

Ploetz, R. C., Hulcr, J., Wingfield, M. J., and De Beer, Z. W. 2013. Destructive tree diseases associated with ambrosia and bark beetles: black swan events in tree pathology? Plant Disease 97:856–872.

R Core Team. 2013. R: A language and environment for statistical computing.

Rabaglia, R. J., Dole, S. A., and Cognato, A. I. 2006. Review of American Xyleborina (Coleoptera: Curculionidae: Scolytinae) occurring north of Mexico, with an illustrated key. Annals of the Entomological Society of America 99:1034–1056.

Reynolds, D. L., and Burke, K. L. 2011. The effect of growth rate, age, and chestnut blight on American chestnut mortality. Castanea 76:129–139.

Roy, B. 1993. Patterns of rust infection as a function of host genetic diversity and host density in natural populations of the apomictic crucifer, *Arabis holboellii*. Evolution 47:111–124.

Seo, M., Martini, X., Rivera, M. J., and Stelinski, L. L. 2017. Flight capacities and diurnal flight patterns of the ambrosia beetles, *Xyleborus glabratus* and *Monarthrum mali* (Coleoptera: Curculionidae). Environmental Entomology 46:729–734.

Shields, J., Jose, S., Freeman, J., Bunyan, M., Celis, G., Hagan, D., Morgan, M., Pieterson, E. C., and Zak, J. 2011. Short-term impacts of laurel wilt on redbay (*Persea borbonia* L. Spreng.) in a mixed evergreen-deciduous forest in northern Florida. Journal of Forestry 109:82–88.

Shimwela, M., Halbert, S. E., Keremane, M., Mears, P., Singer, B., Lee, W. S., Jones, J. B., Ploetz, R., and van Bruggen, A. H. 2019. In-grove spatiotemporal spread of citrus huanglongbing and its psyllid vector in relation to weather. Phytopathology:418–427.

Shimwela, M., Schubert, T., Albritton, M., Halbert, S., Jones, D., Sun, X., Roberts, P., Singer, B., Lee, W., and Jones, J. 2018. Regional spatialtemporal spread of citrus huanglongbing is affected by rain in Florida. Phytopathology 108:1420–1428.

Therneau, T., Atkinson, B., and Ripley, B. 2015. rpart: Recursive partitioning and regression trees. R package version 4:1–9.

Welsh, C., Lewis, K., and Woods, A. 2009. The outbreak history of Dothistroma needle blight: an emerging forest disease in northwestern British Columbia, Canada. Canadian Journal of Forest Research 39:2505–2519.

Wong, C. M., and Daniels, L. D. 2017. Novel forest decline triggered by multiple interactions among climate, an introduced pathogen and bark beetles. Global Change Biology 23:1926–1941.

Wuest, C. E., Harrington, T. C., Fraedrich, S. W., Yun, H.-Y., and Lu, S.-S. 2017. Genetic variation in native populations of the laurel wilt pathogen, *Raffaelea lauricola*, in Taiwan and Japan and the introduced population in the United States. Plant Disease 101:619–628.

Zhou, Y., Avery, P. B., Carrillo, D., Duncan, R. H., Lukowsky, A., Cave, R. D., and Keyhani, N. O. 2018. Identification of the Achilles heels of the laurel wilt pathogen and its beetle vector. Applied Microbiology and Biotechnology 102:5673–5684.

